# Heterogeneity and Intrinsic Variation in Spatial Genome Organization

**DOI:** 10.1101/171801

**Authors:** Elizabeth H. Finn, Gianluca Pegoraro, Hugo B. Brandão, Anne-Laure Valton, Marlies E. Oomen, Job Dekker, Leonid Mirny, Tom Misteli

**Affiliations:** National Cancer Institute, NIH, Bethesda, MD 20892; High-throughput Imaging Facility, National Cancer Institute, NIH, Bethesda, MD 20892; Graduate Program in Biophysics, Harvard University, Cambridge, MA 02138; Howard Hughes Medical Institute, Program in Systems Biology, Department of Biochemistry and Molecular Pharmacology, University of Massachusetts Medical School, Worcester, MA 01605; Institute for Medical Engineering and Science and Department of Physics, Massachusetts Institute of Technology, Cambridge, MA 02139

## Abstract

The genome is hierarchically organized in 3D space and its architecture is altered in differentiation, development and disease. Some of the general principles that determine global 3D genome organization have been established. However, the extent and nature of cell-to-cell and cell-intrinsic variability in genome architecture are poorly characterized. Here, we systematically probe the heterogeneity in genome organization in human fibroblasts by combining high-resolution Hi-C datasets and high-throughput genome imaging. Optical mapping of several hundred genome interaction pairs at the single cell level demonstrates low steady-state frequencies of colocalization in the population and independent behavior of individual alleles in single nuclei. Association frequencies are determined by genomic distance, higher-order chromatin architecture and chromatin environment. These observations reveal extensive variability and heterogeneity in genome organization at the level of single cells and alleles and they demonstrate the coexistence of a broad spectrum of chromatin and genome conformations in a cell population.

---

Genomes exist in a highly organized fashion in the nucleus of human cells (*1, 2*). The folding and looping of DNA gives rise to conserved elements of organization including non-random radial positions of chromosome territories within the 3D nuclear space (*3, 4*), separate compartments for heterochromatin and euchromatin (*5, 6*), < 1 Mbp-scale domains, referred to as topologically associated domains (TADs), promoter-enhancer loops which control gene expression (*7-9*), and functional clusters of gene loci (*10, 11*) such as the ribosomal genes in the nucleolus (*12*). Studies of spatial genome organization have relied extensively on complementary imaging approaches and biochemical methods that capture different aspects of genome organization and may be hard to compare to each other directly (*13, 14*). While microscopy-based methods, such as fluorescence in situ hybridization (FISH), allow direct measurements of three-dimensional distances between loci, they are typically limited to probing of a low number of candidate loci. Biochemical methods, such as high-throughput chromosome conformation capture (Hi-C) or ChIA-PET, on the other hand, provide genome wide maps of interactions frequencies, and hence are most sensitive to small distances that allow capturing of the interaction, but insensitive to larger distances. While providing maps of interaction frequencies for the entire genome and at high resolution (*15-17*), chromosome conformation capture is routinely performed on populations of millions of cells generating averaged snapshots of the population (*18*). Single cell Hi-C experiments and polymer-model based simulations of Hi-C data have begun to probe variability in genome organization between individual cells (*19-23*), however, these approaches are often limited to low numbers of cells, may suffer from relatively low read coverage, and do not distinguish potential differential behavior of individual alleles in the same nucleus (*24, 25*). Here we have developed an orthogonal approach based on the combined use of high-resolution Hi-C and high-throughput imaging to map the spatial position and colocalization frequencies of hundreds of genomic loci to systematically probe the heterogeneity of physical genome interactions and allele-specific behavior at the single cell level. Our findings reveal a remarkably high degree of cell-to-cell and allele-to-allele heterogeneity of spatial genome organization.

To probe the patterns and extent of variation in genome organization in a cell population, we applied High-throughput Imaging Position MAPing (HIPMap; *26*) based on high-throughput fluorescence in situ hybridization (hiFISH) to systematically determine the spatial position and distances between combinations of genomic interaction pairs identified by Hi-C in human foreskin fibroblasts (see Methods; 250kb resolution used for probe finding). A test set of 91 Hi-C site pairs on chromosomes 1, 17, and 18 (Fig 1A) were selected so as to maximize the range of Hi-C frequency (500-fold range), to provide sets of distance-matched sites (similar genomic distances, up to 10-fold range in Hi-C frequency), Hi-C-matched sites (similar Hi-C frequency, genomic distance ranging from 10 Mb to 250 Mb), and to represent gene rich as well as gene poor regions (Fig 1A). In addition, two genome regions of 2.88 and 2.75 Mb, respectively, on chromosome 4 containing multiple TADs defined by Hi-C were tiled at higher density to examine short range associations (Fig 1A, see Methods). Bacterial Artificial Chromosome (BAC) probes were selected for each region of interest, and corrected Hi-C maps and bias vectors were calculated using 250kb bins centered at their midpoint (*27*). Combined, a total of 136 site pairs were initially tested by FISH. Twenty of these pairs showed either Hi-C bias scores in the top 5% or were flagged as outliers in the distribution of Hi-C values (i.e. with z-scores over 2 compared to all equidistant pairs; coverage maps in Fig S2). Over 1000 minimum physical distance measurements of FISH signals were determined for each locus pair in 2D and 3D using an optimized image analysis pipeline (*28*). Trends were similar using both 2D and 3D distances, and to maximize accuracy 3D distances were used throughout. The distribution of minimum distances between loci showed little overlap with the distribution of maximum distances, especially for proximal regions (Fig S1A), demonstrating that this approach differentiates pairs in *cis*, on the same chromosome, from those in *trans*, between alleles. Distance measurements were not affected by chemical fixation and processing during the hiFISH procedure and reflected in vivo distances as demonstrated by comparable distance distributions between LacO and TetO portions of a previously described 25kb LacO-TetO array in live cell imaging of NIH 2/4 mouse fibroblasts before and after fixation and permeabilization (Fig S1B, C) (*29*).

**Fig. 1.**
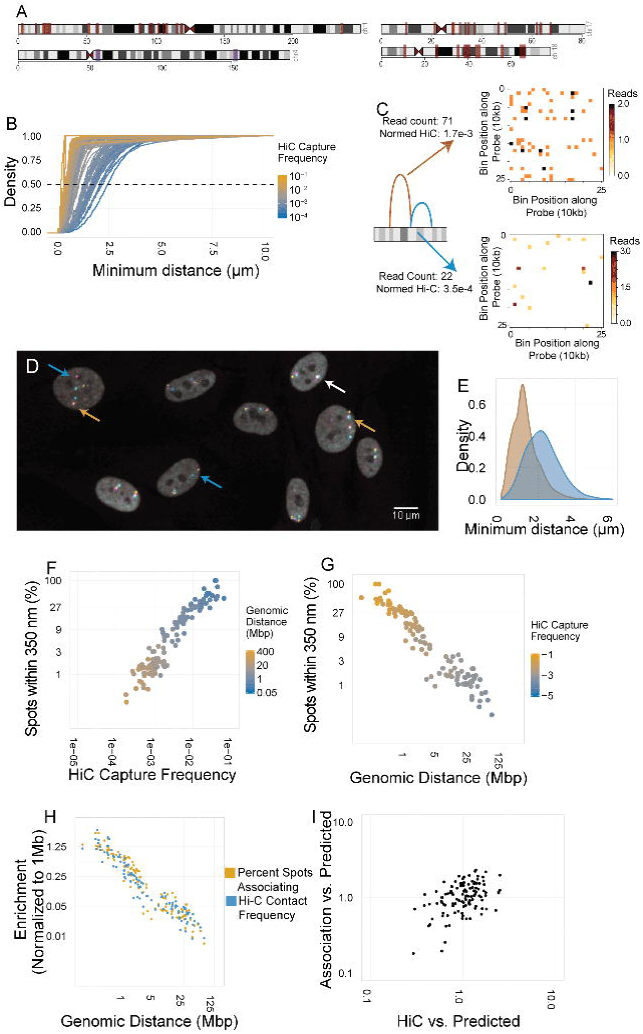
Spatial mapping of genome interactors. (**A**) Ideogram of a set of interaction partners on chromosomes 1, 4, 17, and 18 used for spatial mapping of genome interactions. Sites are indicated by red bars, most pairwise combinations of sites within a chromosome were tested. (**B**) Cumulative distance distributions showing the range of minimal distances for all tested pairs. Median (50% total density) indicated by dashed line. Color indicates Hi-C capture frequency (blue: low, orange: high). (**C**) Schematic diagram of “interactor” and “non-interactor” tested pairs, summed read counts throughout the entire 250 kb bin are shown with coverage plots showing reads per 10kb bin throughout the 250 kb bin. (**D**) Representative FISH image showing a field of cells stained for three loci on chromosome 1. Yellow: bait probe (Chr1 LR.52, chromosome 1, 12,768,721- 12,925,598); Cyan: “non-interacting” target probe (Chr1 LR.91, chromosome 1, 22,549,855- 22,721,150); Magenta: “interacting” target probe (Chr 1 LR.11, chromosome 1, 2,301,890- 2,502,158). Arrangements where all sites were separated by large distances indicated by blue arrows, arrangements where all sites colocalize indicated by white arrow, arrangements where one but not both sites colocalize indicated by orange arrows. (**E**) 3D distance distributions showing the range of minimal distances between “interactor” pair (brown) and “non-interactor” pair (blue). (**F**) Scatter plot showing the percentage of spot pairs within 350 nm vs. Hi-C capture frequency. Color indicates genomic distance. (**G**) Scatter plot showing the percentage of spot pairs within 350 nm vs. genomic distance; color indicates Hi-C capture frequency. (**H**) Scatterplot showing enrichment of percent spots associating or Hi-C capture frequency vs. 1 Mb average as for each site pair vs. genomic distance. Percent spots associating: orange, Hi-C capture frequency: blue. (**I**) Scatterplot of values normalized by distance-based predictions. Predicted values were generated based on a log:log model, and the ratio between predicted and observed value was used. X-axis: Hi-C capture frequency/prediction based on genomic distance. Y-axis: Percent spots associating/prediction based on genomic distance.

To establish how Hi-C interaction frequencies relate to physical distances, we first calculated cumulative distributions of pairwise spatial distance for each site in the entire set (Fig 1B). The median spatial distances of site pairs strongly correlated with their Hi-C frequencies (Fig 1B). Importantly, the distance distributions lacked multimodality or discontinuity, suggesting that the overall population of distances is variable and continuous, but in any individual cell non-interactors may often be found at shorter separations than interactors, consistent with other findings for a few specific loci (21,24). This conclusion was corroborated by analysis of the distance distribution of a single bait probe on chromosome 1 and either an 10Mb upstream Hi-C “interactor” (71 total reads, normalized Hi-C capture frequency of 1.7*10^−3^) or an equidistant downstream “non-interactor” (22 total reads, normalized Hi-C capture frequency of 3.5*10^−4^; Fig 1C, representative image 1D). Despite bias scores of these probes in the top 5% suggesting poorly mappable regions with unreliable Hi-C frequencies (Fig. 1C), we observe a statistically significant difference in 3D spatial distance between these two targets with the “interactor” at an average distance of 1.30 µm from the bait and the “non-interactor” at an average distance of 2.26 µm (Fig 1E, p<2.2*10^−16^). However, the distributions strongly overlap, with an overlapping coefficient of 0.57, and 22% of bait spots were physically closer to the the “non-interactor” target than the “interactor” target (Fig 1E). Similar results were observed with 5 additional distance-matched pairs of one bait and two targets, all showing even larger overlaps, and all distance distributions at all genomic distances were highly overlapping (Fig 1B). Furthermore, analyzing the noise at all 91 long-range locus pairs via both Fano factor (variance in distance/mean distance) and coefficient of variation (variance in distance/mean distance squared) shows that while variation moderately correlates with mean spatial distance, the most variable regions tend to be close together in the genome (< 1 Mb) and display high Hi-C capture frequencies, demonstrating that variability is greatest at sites that are most likely to interact (Fig S3). These observations demonstrate correlation of physical distance with Hi-C data, but at the same time reveal extensive heterogeneity in physical proximity of Hi-C interaction pairs in individual cells.

To more directly relate physical distance to Hi-C interaction frequency (*13*), and to determine the prevalence of a given interaction in a cell population, we measured the colocalization frequencies of interaction pairs in the population (Fig 1G, H). We find strong correlations between Hi-C frequency and spatial proximity for all pairs at 3D distance thresholds of 150 nm, 200 nm, 350 nm and 1um (Fig 1F, Fig S4). The percentage of pairs within 350 nm for the most common interactors were approximately 200-fold enriched relative to the least common interactors (0.25% to 53.4% of alleles; r^2^ = 0.904, p < 2.2*10^−16^). Similar trends were observed upon relaxing the distance threshold to 1 µm (r^2^ = 0.829, p < 2.2*10^−16^; 40-fold change from 2.42% to 99.5% of alleles, Fig S4), though showing saturation for large interactions frequencies, and reduction of the threshold to 150 nm (r^2^: 0.861, p < 2.2*10^−16^; ~750-fold change from 0.04% to 27.3% of alleles, Fig S4). The similarity of trends at different thresholds demonstrates that the observed results are not significantly influenced by microscope drift or optical aberrations. Of note, the percentage of alleles engaged in a physical interaction at any given time in the population is low. Most probed interaction pairs (109/146) showed colocalization within 350 nm in fewer than 30% of alleles, and few interaction pairs (16/146) showed colocalization in more than 50% of alleles (Fig 1F). At 150 nm, most pairs (132/146) colocalized less than 10% of the time, and only very few (4/146) colocalized more than 15% of the time. The low frequency of physical association provides a rationale for the observed continuum of spatial distances and the lack of multimodality in the distance distributions (Fig 1F). Analysis of residuals using a power-law model indicated a typically ~2.6-fold range in percentage of colocalizing signals for most pairs of similar Hi-C capture frequency, for example distal site pairs with a ±10% range in Hi-C frequencies associate within 350 nm between 1 and 3% of the time and within 150 nm 0.1 and 0.6% of the time; more proximal pairs with a ±10% range colocalize between 16 and 40% of the time for separation within of 350, and 2 and 9% within 150 nm (Fig S4). These results demonstrate overall low frequency of Hi-C interactions and a remarkably high degree of variability in association frequencies and spatial separation of genome regions in individual cells across a population.

The most prominent contributor to the likelihood of physical association of two loci is the genomic distance separating them. We find a strong inverse correlation between association frequency and genomic distance (Fig 1G; at 350 nm, r^2^ = 0.907, p < 2.2*10^−16^; at 1 µm, r^2^ = 0.877, p < 2.2*10^−16^). As observed for Hi-C frequencies, root mean squared error (RMSE) indicated typically a two to three-fold variation in percent colocalizing for locus pairs at similar genomic distances; for example, two locus pairs separated by very similar genomic distances (412 and 419 kb) were found to colocalize with significantly different frequencies of 46% and 26% of alleles, respectively (Fig 1G). The genomic distance-related decay of the percentage of colocalizing spots is very similar to that of Hi-C capture frequency, indicating a well-known strong contribution of genomic distance to association frequency measured by Hi-C and FISH (Fig 1H). Upon stratification of Hi-C and 350nm colocalization frequency by genomic distance (based on power-law distance-based expected values) weak correlation between Hi-C and colocalization frequency is maintained (r^2^ = 0.138, p = 2.1*10-5, Fig 1I). As a control, the correlation between Hi-C frequency and 350nm association frequency was decreased when Hi-C frequency was replaced with that of random pairs shifted by 500kb were used (r^2^ = 0.801, p < 2.2*10^−16^; Fig S4). These findings suggest that while genomic distance is a major determinant of proximity between two genomic loci, locus-specific properties of the chromatin, such as the formation of loops, TADs or compartments also contribute to the interaction frequencies measured by Hi-C and the colocalization frequencies measured by FISH (see below).

To investigate how chromatin context affects the likelihood of spatial proximity, the 200 loci on chromosomes 1, 17, and 18 were classified by gene density (see Methods). When 350 nm colocalization frequencies were determined, locus pairs between gene rich regions showed a weaker correlation with genomic distance than loci in gene poor regions (Fig 2A; gene poor: r^2^ = 0.929 and p < 2.2*10^−16^; gene rich: r^2^ = 0.811 and p < 2.2*10^−16^). Gene rich loci also exhibited a wider range of association frequencies than gene poor regions (Fig 2B). This trend was significant on chromosome 1 between gene rich and gene poor regions (F-test, p=0.00781, ratio between variances: 0.513) as well as between chromosome 17 and chromosome 18 (F-test, p= 0.0286, ratio between variances: 0.45), which have about the same size but 6-fold difference in the number of genes (1200 in chr 17, and 270 in chr 18). These observations suggest greater distance dependence and smaller variation of physical association between locus pairs in gene poor regions of the genome than those in gene rich regions. This conclusion is consistent with a model in which gene-poor chromatin is denser and characterized by nonspecific interactions resulting in chromatin compaction, whereas gene-rich chromatin is overall more dispersed, but prone to specific interactions such as locus-specific loops (*6*). Furthermore, classification of site pairs according to their localization to the A or B compartments of chromatin (see Methods), which generally correlate with chromatin state, demonstrated associations between loci occur with slightly higher frequency among pairs where both loci belong to compartment B than A-A or A-B pairs (Fig 2D). This is consistent with a well-known compartmentalization of the genome observed in Hi-C data where BB and AA interactions are about 2-fold more frequent than AB interactions, as seen here in observed-over-expected Hi-C (Fig 2C). This behavior is consistent with the conceptual model of A and B compartments representing distinct sets of interactors (*15*), and with the notion that B compartments are more compact and self-associating, while A compartments are more dispersed and interact more uniformly with all genomic loci (*6,15*).

**Fig. 2.**
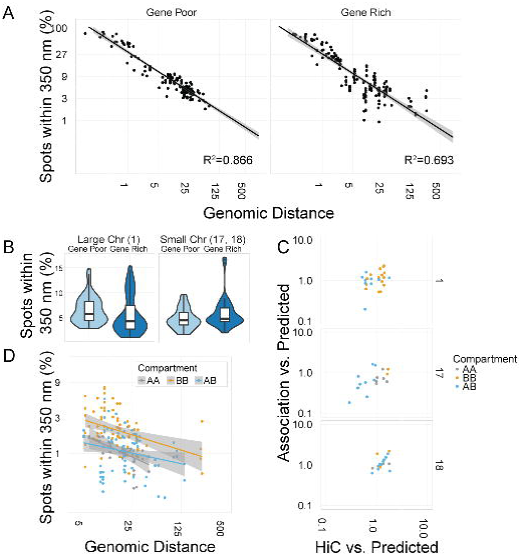
Modifiers of association frequencies. (**A**) Scatterplots showing the percentage of spot pairs within 350 nm vs. genomic distance, separated into gene poor or gene rich site pairs. Line of best fit and R^2^ values indicated on plots. (**B**) Box/violin plots showing total range in percentage of spot pairs within 350 nm for all pairs tested, by chromosome and gene content. (**C**) Scatterplots showing model-normalized association likelihood vs Hi-C score for each chromosome, color-coded by compartment. Gray: both sites in compartment A; orange: both sites in compartment B; blue: one site in each compartment. (**D**) Scatterplot showing percentage of spot pairs within 350 nm vs. genomic distance, color-coded by compartment of sites.

TADs are regions of the genome characterized by elevated interaction frequencies for locus pairs within the region as compared to locus pairs at similar distances outside the region (*15*). To probe the variability of spatial organization of TADs in a population, we selected two regions on chromosome 4 encompassing a total of 14 TADs and measured distances at locus pairs within the same TAD at varying distances (from 70 kb to 600 kb) and across an 8-fold range of Hi-C frequencies (Fig 3A). As expected, the frequency of association of locus pairs within the same TAD was high with 57 to 84% of alleles showing colocalization within 350 nm (Fig 3B, green spots), mostly because of their smaller genomic separation. We observe limited correlation between genomic distance and percent spots associating within 350 nm or 150 nm within the same TAD (350 nm: r^2^ = 0.107, p = 0.135; 150 nm: r^2^ = 0.086, p = 0.162; Fig 3B, top panels) nor correlation between Hi-C frequency and percent spots associating within the same TAD (350 nm: r^2^ = 0.129, p = 0.113; 150 nm: r^2^ = 0.132, p = 0.110; Fig 3B, bottom panels). Considering edge-to-edge rather than center-to-center distances did not recover correlations within a TAD (350 nm: r^2^ = 0.196, p = 0.064; 150 nm: r^2^ = 0.040, p = 0.246; Fig S5A). The absence of correlation is not due to limited resolution, microscope drift or optical aberration, as we can robustly differentiate a single locus stained in two colors from two loci within the same TAD (Fig S5B, C). However, it is possible that these correlations are lost due to the fact that BAC probes are on a similar length-scale to many TADs, and frequently cross TAD borders. In contrast to intra-TAD interactors, loci in adjacent TADs separated by larger distances from 250 kb to 1 Mb and a 4-fold range in Hi-C capture frequency colocalize only slightly less often than those within TADs (40-72%), and they showed an increase in correlation between the percentage of alleles within 350 nm and either Hi-C or genomic distance (Hi-C; r^2^ = 0.204 and p = 0.039; genomic distance r^2^ = 0.320 and p = 0.011; Fig 3B). The distinct correlation behavior of intra- vs. inter-TAD interaction pairs is in line with the interpretation that although TADs are distinct domains as reflected in Hi-C data (*30*), new models and single-cell Hi-C suggest that any specific TAD is a highly dynamic and variable domain, overlapping in space with its neighbors and is a reflection of the population average of enriched, but highly variable, interactions between individual sites (*23,31,32*).

**Fig. 3:**
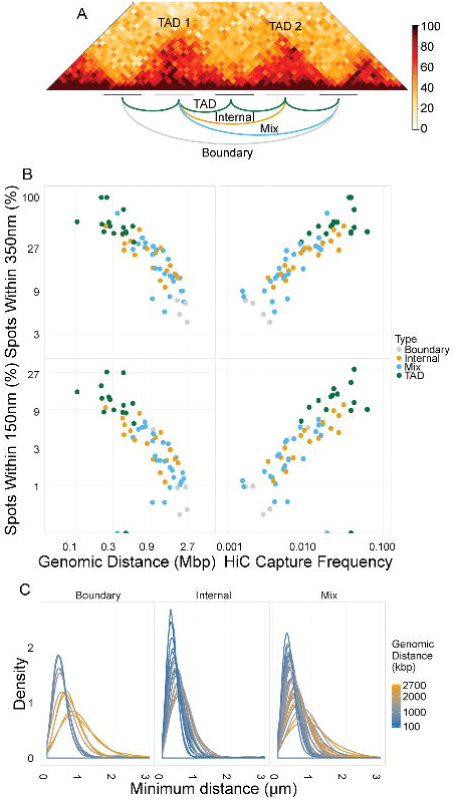
Intra- and Inter-TAD interactions. (**A**) Diagram showing probe tiling arrangement, with representative Hi-C heatmap and example TAD, Internal, Mix, and Boundary pairs marked. (**B**) Scatterplots showing the percentage of spot pairs within 350 nm (top panels) or 150 nm (bottom panels) versus either genomic distance (left panels) or Hi-C capture frequency (right panels). (**C**) Distance distributions showing the range of minimal distances for each site pair (line). Color-coded by genomic distance.

One common hypothesis for TAD organization is their arrangement in “looped florets”, in which the boundary sequences between TADs are brought together, possibly by architectural proteins (*30, 31*). A prediction from this model is that the bases of multiple TADs are in closer spatial proximity than two regions in the interior of neighboring TADs. To test this model, we analyzed distance distributions between multiple TAD boundaries (“boundary” pairs), between internal regions in different TADs (“internal” pairs), or between a TAD boundary and a region internal to another TAD (“mixed” pairs) (Fig 3A). Scatterplots comparing the percent of colocalization within 350 nm relative to genomic distance showed internal, boundary, and mixed pairs all to be roughly collinear (Fig 3B). Fitting with a power-law model revealed that boundary pairs were the most distance dependent (r^2^ = 0.89, scaling factor = −1.75 ± 0.01), followed by mixed pairs (r^2^ = 0.59, scaling factor = −0.95 ± 4.12*10^−6^) and internal pairs (r^2^ = 0.70, scaling factor = −0.59 ± 3.39*10^−5^). Considering only site pairs separated by at least 750 kb or fitting an exponential rather than a power-law curve to the data did not change the trends or r^2^ values. These differences in scaling suggest that internal regions, rather than TAD boundaries, are more likely to interact between TADs. Furthermore, the distributions of all types of pairs, matched for genomic distance, were comparable, corroborating the absence of strong boundary clustering (Fig 3C; color coded by genomic distance). Finally, while a “looped floret” model predicts that clusters of three or more boundary elements would be common, we do not find this to be the case: no tested site triplet of regions in neighboring TADs occurred more than 25% of the time, and no triplet occurred significantly more or less than expected based on pairwise association frequencies (Fig S5D), consistent with independent looping interactions (31). The observations suggest that TAD boundaries, although they form the edges of internally interacting regions, are not universally themselves the bases of stable loops, consistent with the model where such loops are dynamically formed by loop extrusion and dissolved (*32, 33*).

Our observations demonstrate considerable variance in spatial distance between any two loci, regardless of their likelihood of interaction. The variability measured in the population may represent heterogeneity in the behavior of alleles in individual cells, or it could be the result of cell-to-cell variation while the behavior of alleles in the same cell is coordinated. To directly measure the cell-specific variability of association frequencies, we compared for each of the 91 long-range locus pairs the behavior of the two alleles in the same nucleus by measuring minimal distances of both alleles relative to their interaction partners (Fig 4A). Scatter plots representing the minimal distance at one allele vs. the minimal distance at the second allele in each nucleus (selected plots, Fig 4A; all plots Fig S6A) demonstrated little correlation between the behavior of the two alleles for any of the 91 locus pairs (Fig 4A). When individual nuclei were categorized based on how many colocalization events they contained (0, 1, or 2), the vast majority of cells contained no events (70.04-99.46%), a limited fraction (0.54-24.24%) contained one event, and very few (0.02-5.34%) of cells contained two events (Fig 4B), in line with the observed low prevalence of overall interactions in the population (Fig 1). These values were generally only slightly enriched relative to expected values calculated based on pairwise association frequencies under the assumption of independence between the two associations (Fig 4B). Enrichment beyond the expectation value for cells with two associations was observed at a few loci, including the most common interactors (selected: Fig 4B; all Fig S6B), however, in no case did the population of cells containing two interactions represent more than 3% of cells (Fig 4B). These data reinforce the observed low prevalence of colocalization throughout the population and demonstrate that interactions most frequently occur independently at the two homologous chromosomes within a nucleus.

**Fig. 4:**
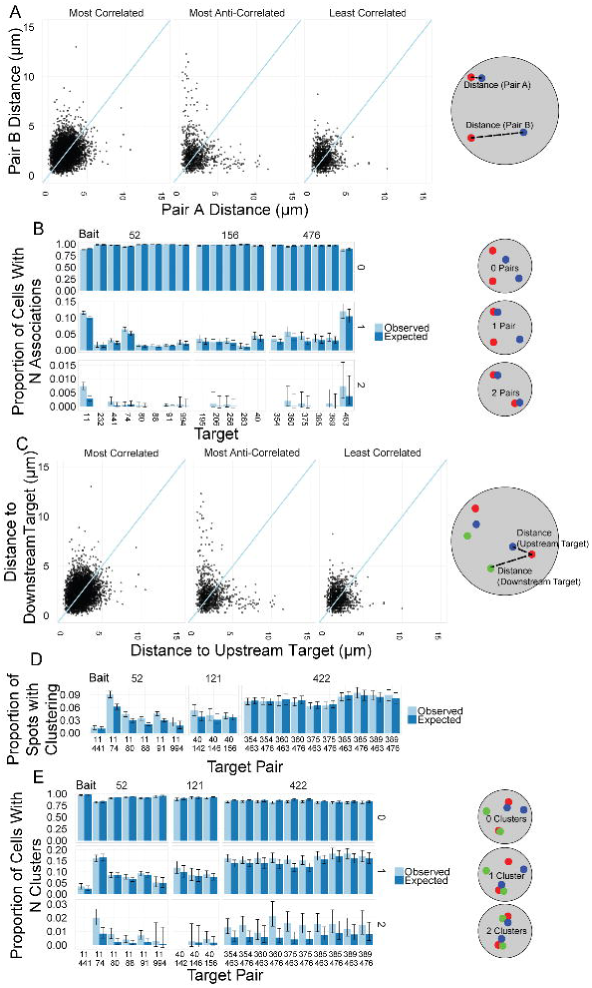
Allelic independence and clustering. (**A**) Scatterplots of minimal distances of both alleles of an interaction pair on a per-cell basis at the most correlated site pair, least correlated site pair, and most anti-correlated site pair (as determined by slope of line and r^2^; distances as diagrammed at right). (**B**) Bar graph showing proportion of cells with 0, 1, or 2 associations within 350nm between spots for a variety of selected site pairs (as diagrammed at right). Expected values calculated based on pairwise spot association probabilities are indicated for reference (see methods). (**C**) Scatterplots showing correlation of distances between bait and two targets (upstream target and downstream target) on a per-bait basis, for most correlated, most anti-correlated, and least correlated triplets (as determined by slope of line and r^2^; diagrammed on right). (**D**) Bar graph showing proportion of bait spots with a triplet association within 350 nm for selected triplets. Expected values calculated from pairwise spot association probabilities are indicated for reference (see methods). (**E**) Bar graph showing proportion of cells with 0, 1, or 2 triplet associations for selected triplets (as diagrammed on right). Expected values calculated from pairwise spot association probabilities are indicated for reference (see Methods).

In addition to pairwise associations, multiple gene loci may also physically form clusters, possibly due to mutual stabilization of pairwise interactions and possibly dependent on chromatin context (*31, 34*). To globally assess the extent and cell-to-cell variability of clustering of multiple interaction pairs, we measured the degree of covariation between multiple site pairs on the same chromosome and generated scatter plots indicating the spatial distance from a single bait to a first target vs. the distance from the same bait to a second target (Fig 4C). Analysis of probes consisting of 18 different baits combined with 18 different downstream targets and 12 different upstream targets representing 43 triplets within the sampled loci on chromosome 1, chromosome 17, and chromosome 18, revealed little evidence of clustering of multiple pairs, as indicated by a lack of accumulation of points along the diagonal, with a maximum r^2^ = 0.1311 for all combinations (select examples Fig 4C; all Fig S7A). Enrichment over expected values as calculated from pairwise association frequencies was detected sporadically, for instance for probes 375/354/422 or 11/52/91 in which triplets were observed 1.67 times (r^2^ = 0.1162) and 1.5 times (r^2^ = 0.04929) more often than expected, respectively (Fig 4D). However, cluster formation was overall rare and occurred at less than 15% of loci for even the most enriched clusters. These results were confirmed by direct counting of clusters in individual cells with 66-99% containing no triplet clusters, <30% containing one cluster <5% containing two clusters (Fig 4E). While Hi-C data is not well suited to detect triplet interactions (*35, 36*), our observations suggest that non-specific clustering of multiple interaction pairs is not a general principle of higher order genome organization, suggesting that reported clustering of genes with complex regulatory hubs (*7, 8, 34, 37, 38*) may be a locus-specific rather than general effect, or may reflect their tendency to interact as seen in population-averaged data rather than formation of such hubs in individual cells.

We demonstrate here a remarkably high level of heterogeneity in spatial genome organization amongst individual cells and between alleles in the same cell. Our conclusions are based on the combined used of Hi-C datasets and high-throughput imaging to determine the variability of genome architecture at the single cell level. This approach takes advantage of the genome-wide nature of Hi-C datasets and the ability of high-throughput imaging to probe genome organization at the single cell level and with high statistical power due to imaging of a large set of locus pairs in a large number of individual cells. The method overcomes the limitations of traditional biochemical mapping methods which generate population averaged datasets and at the same time elevates traditional imaging methods beyond their anecdotal nature due to their small sample sizes. Our observations complement single cell Hi-C analyses and extend them by probing the behavior of individual alleles in thousands of nuclei.

We find a remarkable level of variability in how the genome folds in individual cells. Long range associations (> 5 Mb) typically occur in no more than 10% of cells for any particular locus pair, while a small number of such distal associations are present in up to 20-25% of cells. As expected, loci within a TAD colocalize at higher frequencies (30-80%). The relatively low frequency of mid- and long range range associations in the population and the combinatorial occurrence of multiple associations in individual cells generates a wide spectrum of genome-wide organizational patterns in the cell population. The observed variability in spatial proximity seems not to be due to extrinsic factors such as signaling or the presence of stable subpopulations of functionally distinct cells, since in that case coordination of alleles would be expected. On the contrary, we observe little correlation between the two alleles in a single nucleus and we find that the two pairs within a single nucleus generally behave independently of each other. This property suggests instead that much of the observed variability does not reflect cell-autonomous genome organization but instead represents intrinsic variability within individual genomes. It is possible that the observed lack of colocalization of interactors is a reflection of dynamic cycles of association and dissociation in the living cells. This explanation, however, seems unlikely, since live cell imaging has demonstrated the motion of DNA in eukaryotic cells to be generally constrained within a radius of less than 1.5 µm (*39-41*), whereas even the most common long-range interactors mapped here are often found at significantly larger distances that can not be attributed to intrinsic dynamic motion. Such high variation in genomic interactions and spatial distances can be well-explained by the polymer nature of chromatin. Consistent with the central role of the polymer nature of chromatin is strong dependence of mean spatial distances and association frequencies on genomic separation between loci.

While genomic distance accounts for most of the variation in interaction probability, with more proximal loci being more likely to interact, our data point to several other factors as determinants of interaction frequencies, including gene density since loci with more genes tend to show less distance dependence and a greater likelihood for very long range (>10 Mb) interactions, which in turn agrees with Hi-C interactions frequencies that depend on compartment type. We also find a contribution of the chromatin context since loci in the A compartment are overall less likely to interact and show different scaling according to distance compared to those in the B compartment.

Our data provide direct experimental support for conclusions from early predictive computational models based on tethered conformation capture (TCC) which suggested that interaction maps are the composite of multiple genome configurations and that the patterns of individual interactions differ between cells (*42*). This interpretation is consistent with our results that any given spatial colocalization at mid- and long-range genomic distances (> 5 Mb) occurs in only 3 to 25% of cells. Furthermore, recent single cell Hi-C studies have demonstrated that, while interactions most commonly occur within a TAD or between TADs of the same compartment, individual cells exhibited widely distinct TADs and some interactions between compartments (*21-23, 43*). Our analysis of thousands of individual cells highlights that associations between loci within the same TAD are the only pairing events present in a large fraction of cells, but notably not all, whereas associations between adjacent TADs or compartments are highly variable giving rise to the observed distinct TAD patterns in single cell Hi-C data. Taken together, our data determine in a systematic and quantitative fashion the extent and nature of the heterogeneity and variability in higher order genome architecture. Our results support a view in which genome organization is remarkably plastic, heterogeneous and marked by variability between individual alleles and cells. How functional interactions can be established despite such highly variable organization is a challenging question that will need to be addressed by future research.

## Materials and Methods

### Cell Culture

Human foreskin fibroblasts (HFF) immortalized with hTert via neomycin selection as described previously (*44*) were grown in DMEM with 10% FBS, 2mM glutamine, and penicillin/streptomycin and split 1:4 twice weekly. Their karyotype was verified as normal via SKY staining (data not shown). HFFs were plated at a density between 5000 and 7500 cells per well in 384 well plates (CellCarrier or CellCarrier Ultra, Perkin Elmer), and left to grow and settle overnight before fixation in 4% paraformaldehyde for 10 minutes. After fixation, plates were rinsed twice in PBS and stored in 70% ethanol at −20°C for up to two months. A total of 40 hiFISH experiments were performed in 42 wells and 14 probe combinations (3 wells per probe combination) per experiment. 500-1000 cells were imaged per probe combination per experiment.

### High-throughput Chromosome Conformation Capture

Hi-C libraries were generated from HFF cells cross-linked in 1% formaldehyde as described previously (*15, 45, 46*). Four technical replicates of 25 million cells each were generated. The libraries were sequenced using 50bp paired-end reads with a HiSeq2000 machine with each replicate in its own lane and replicates 3 and 4 sequenced two more times in one lane. The eight lanes were pooled; the total number of valid pairs after mapping and filtering was 746,195,659. The filtered reads were mapped to the human genome (hg19) using Bowtie 2.2.8 and normalized using the iterative mapping strategy described previously (*47, 48*).

### Probe selection

For long-range interactions (> 10Mb separation) within chromosomes, the single-chromosome Hi-C map with 250kb binning was used and associations classified into proximal (separated by ~ 10 Mb or 10% of the chromosome), medial (separated by ~ 50 Mb or 30% of the chromosome), and distal (separated by ~ 100 Mb or 50% of the chromosome) and subsequently ranked into likely interactors (top 100 interactions based on Hi-C signal at a given distance), unlikely interactors (bottom 100 interactions at a given distance), or median (median 100 interactions at a given distance). Probe families were selected as those with a single bait and representative targets from as many categories (e.g. proximal, likely interactors vs. distal, likely interactors) as possible.

For short range interactions (< 3 Mb separation), two regions on chromosome 4 were chosen by visual inspection to contain several adjacent TADs likely within the same compartment. These regions were tiled using adjacent/overlapping BAC probes, using, if possible, probes to position a BAC probe at the base/boundary between each TAD-like structure and one within each TAD-like structure. For TADs that were too small for tiling without substantial overlap between adjacent BAC probes, only boundary or only interior probes were chosen.

A table of the start and end locations of all BACs used is included as Table S1; a table of all pairwise associations tested, along with recalculated Hi-C scores in the centered 250 kb bin, bias scores, and percent spots associating is included as Table S2.

### HiFISH Imaging

High-throughput fluorescence in-situ hybridization (hiFISH) was performed as described previously (*49, 50*). Probes were generated via nick translation as described previously (*51*) from bacterial artificial chromosomes (BACs) to approximately 100 genomic loci (see Table 1). Mixes, reagents, and conditions are exactly as in Finn (*28*). All experiments were performed in triplicate.

Imaging between chromosomes and at long distances was performed in four channels (405, 488, 561, 640 nm excitation lasers) in an automated fashion using a dual spinning disk high-throughput confocal microscope (PerkinElmer Opera QEHS) using a 40X water immersion lens (NA = 0.9) and pixel binning of 2 (pixel size = 320 nm). 20-40 fields of view were imaged per well. In all exposures the light path included a primary excitation dichroic (405/488/561/640 nm), a 1^st^ emission dichroic longpass mirror: 650/ 660- 780, HR 400-640 nm and a secondary emission dichroic shortpass mirror: 568/ HT 400- 550, HR 620-790 nm. In exposure 1, samples were excited with the 405 and 640 nm lasers, and the emitted signal was detected by two separate 1.3 Mp CCD cameras (Detection filters: bandpass 450/50 nm and 690/70 nm, respectively). In exposure 2, samples were excited with the 488 nm laser and the emitted light was detected through a 1.3 Mp CCD camera (Detection filter: bandpass 520/35). In exposure 3, samples were excited with the 561 nm laser and the emitted light was detected through a 1.3 Mp CCD camera (Detection filter: bandpass 600/40). Samples were optically sectioned in z every 1 μm for a final volume of 7 μm. For clustering experiments, samples were optically sectioned in z every 300 nm for a final volume of 7 μm. For short range interactions on chromosome 4, imaging was performed in four channels (405, 488, 561, and 640 nm excitation lasers) in an automated fashion using a dual spinning disk high-throughput confocal microscope (Yokogawa CV7000), using a 60x water immersion lens (NA = 1.2) and no pixel binning (pixel size = 108 nm). 16 fields of view were imaged per well. In exposure 1, samples were excited with the 405 and 561 nm lasers, and the emitted light was collected through a path including a short pass emission dichroic mirror (568 nm) and two sCMOS cameras (5.5 Mp) in front of 445/45 nm and 600/37 nm bandpass emission filters, respectively. In exposure 2, samples were excited with the 488 and 640 nm lasers, and the emitted light was collected through the same light path and with the same cameras as exposure 1, but with 525/50 nm and 676/29 nm bandpass emission filters, respectively, instead. Samples were optically sectioned in z every 1 μm for a final volume of 7 μm.

### 2D and 3D Image Analysis

Automated analysis of all images was performed based on a modified version of a previously described Acapella 2.6 (PerkinElmer) custom script (*28, 52, 53*). This custom script performed automated nucleus detection based on the maximal projection of the DAPI image (ex. 405 nm) to identify cells. Spots within these cells were subsequently identified in maximal projections of the Green (ex 488 nm), Red (ex. 561 nm) and Far Red (ex. 640 nm) images, using local (relative to the surrounding pixels) and global (relative to the entire nucleus) contrast filters. The x and y coordinates of the brightest pixel in each spot were calculated. The z coordinate of the spot center was then calculated by identifying the slice in the z-stack with the highest value in fluorescence intensity for each of the spot centers. Datasets containing x,y and z coordinates for spots in the Green, Red and Far Red channels as well as experiment, row, column, field, cell, and spot indices, were exported from Acapella as tab separated tabular text files. These coordinates datasets were imported in R (*52*). 2D and 3D distances for each pair of Red:Green, Red:Far Red, or Green:Far Red probes within a cell were generated on a per-spot basis using the SpatialTools R package (*53*). Subsequent analyses were performed in R using the plyr (*54*), dplyr (*55*), ggplot2 (*56*), data.table (*57*), knitr (*58*) and stringr (*59*) packages. All images, scripts, and datasets are available upon request.

### Statistical Analysis

2D and 3D distances were calculated on a per-green-spot basis using the SpatialTools R package (*53*). All analyses were performed using both 2D and 3D distances as described previously (*28*); 3D distances are reported. For density plots, minimum spot distances were graphed using ggplot2. For scatter plots showing percent associations versus Hi-C capture frequency or genomic distance, as well as box plots showing range in percent spots associating, percent spots colocalizing within a threshold and percent of cells with at least one association were calculated with tools from data.table, and these values were plotted with ggplot2. For scatter plots showing correlations between homologous pairs in the same cell, minimum spot:spot distances were indexed by cell and merged. Pair A and Pair B were assigned arbitrarily. For scatter plots showing correlations between two targets and the same bait, spot pairs were indexed by “central” spot (defined as the middle of three loci as arrayed in the genome) and distances merged. Overlapping coefficient was calculated as the sum of the minimum proportion over all distances: x = Σ_*x*_ min (*p*(*A* ≥ *x* & *A* < *x* + *c*), *p*(*B* ≥ *x* & *B* < *x* + *c*)) where A is the 3D distance between pair A, and B is the 3D distance between pair B, with a step of δ.

## Acknowledgments

We would like to thank Bryan Lajoie for initial mapping and Hakan Ozadam for assistance in organizing and handling Hi-C mapped-pairs files. High-throughput imaging was performed at the High-Throughput Imaging Facility (HiTIF)/Center for Cancer Research, National Cancer Institute, NIH. This research was supported by funding from the Intramural Research Program of the National Institutes of Health (NIH), National Cancer Institute, and Center for Cancer Research and by the 4D Nucleome Common Fund (5U54DK107980-01). H.B.B received support from a Natural Sciences and Engineering Research Council of Canada PGS-D fellowship. J.D. is an investigator of the Howard Hughes Medical Institute.

## Author Contributions

E.H.F. identified pairs, performed hiFISH experiments, and processed hiFISH data. G.P. and E.H.F. developed software for image and statistical analysis. A-L.V. and M.E.O. generated the Hi-C libraries. E.H.F., and H.B.B., performed bioinformatic analysis of Hi-C data and comparisons to hiFISH. E.H.F. and T.M. designed experiments and wrote the manuscript with input from L.M., J.D., H.B.B., and A-L.V.

## References and Notes

1. T. Misteli, Beyond the Sequence: Cellular Organization of Genome Function. Cell 128, 787–800 (2007).

2. W. A. Bickmore, The spatial organization of the human genome. Annual Review of Genomics and Human Genetics 14, 67–84 (2013).

3. T. Cremer et al., Chromosome territories - a functional nuclear landscape. Current Opinion in Cell Biology 18, 307–316 (2006).

4. S. Boyle et al., The spatial organization of human chromosomes within the nuclei of normal and emerin-mutant cells. Hum Mol Genet 10, 211–9 (2001).

5. N. J. Francis, R. E. Kingston, C. L. Woodcock, Chromatin compaction by a polycomb group protein complex. Science 306, 1574–1577 (2004).

6. A. N. Boettiger et al., Super-resolution imaging reveals distinct chromatin folding for different epigenetic states. Nature 529, 418–422 (2016).

7. B. Tolhuis, R. J. Palstra, E. Splinter, F. Grosveld, W. De Laat, Looping and interaction between hypersensitive sites in the active b-globin locus. Molecular Cell 10, 1453–1465 (2002).

8. D. Baù et al., The three-dimensional folding of the α-globin gene domain reveals formation of chromatin globules. Nature Structural & Molecular Biology 18, 107–114 (2011).

9. A. Sanyal, B. R. Lajoie, G. Jain, J. Dekker, The long-range interaction landscape of gene promoters. Nature 489, 109–113 (2012).

10. Q. Wang et al., Cajal bodies are linked to genome conformation. Nature Communications 7, 10966 (2016).

11. D. Rieder et al., Co-expressed genes prepositioned in spatial neighborhoods stochastically associate with SC35 speckles and RNA polymerase II factories. Cellular and Molecular Life Sciences 71, 1741–1759 (2013).

12. F.-M. Boisvert, S. van Koningsbruggen, J. Navasques, A. I. Lamond, The multifunctional nucleolus. Nat Rev Mol Cell Biol 8, 574–585 (2007).

13. G. Fudenberg, M. Imakaev, FISH-ing for captured contacts: towards reconciling FISH and 3C. Nat Methods (2017) doi: 10.1038/nmeth.4329

14. L. Giorgetti, E. Heard, Closing the loop: 3C versus DNA FISH. Genome Biology 17, 215 (2016).

15. E. Lieberman-Aiden et al., Comprehensive Mapping of Long-Range Interactions Reveals Folding Principles of the Human Genome. Science 326, 289–293 (2009).

16. F. Jin et al., A high-resolution map of the three-dimensional chromatin interactome in human cells. Nature 503, 290–294 (2013).

17. G. Li et al., ChIA-PET tool for comprehensive chromatin interaction analysis with paired-end tag sequencing. Genome Biology 11, R22 (2010).

18. J. M. O'Sullivan, M. D. Hendy, T. Pichugina, G. C. Wake, J. Langowski, The statistical-mechanics of chromosome conformation capture. Nucleus 4, 390–398 (2013).

19. R. Kalhor, H. Tjong, N. Jayathilaka, F. Alber, L. Chen, Genome architectures revealed by tethered chromosome conformation capture and population-based modeling. Nat Biotechnol 30, 90–98 (2011).

20. H. Tjong et al., Population-based 3D genome structure analysis reveals driving forces in spatial genome organization. Proc. Natl. Acad. Sci. U. S. A. 113, E1663–1672 (2016).

21. T. Nagano et al., Single cell Hi-C reveals cell-to-cell variability in chromosome structure. Nature 502, 59–64 (2013).

22. S. Carstens, M. Nilges, M. Habeck, Inferential Structure Determination of Chromosomes from Single-Cell Hi-C Data. PLoS Comput. Biol. 12, e1005292 (2016).

23. I. M. Flyamer et al., Single-nucleus Hi-C reveals unique chromatin reorganization at oocyte-to-zygote transition. Nature 544 110–114 (2017).

24. V. Ramani et al., Massively multiplex single-cell Hi-C. Nat Methods 14, 263–266 (2017)

25. T. Nagano et al., Cell cycle dynamics of chromosomal organisation at single-cell resolution. Nature 547 61–67 (2017).

26. S. Shachar, G. Pegoraro, T. Misteli, HIPMap: A High-Throughput Imaging Method for Mapping Spatial Gene Positions. Cold Spring Harbor symposia on quantitative biology (2015).

27. M. Imakaev, G. Fudenberg, R. P. McCord, N. Naumova, A. Goloborodko, B. R. Lajoie, J. Dekker, L. A. Mirny, Iterative correction of Hi-C data reveals hallmarks of chromosome organization, Nat Methods 9 999–1003 (2012).

28. E. H. Finn, G. Pegoraro, S. Shachar, T. Misteli, Comparative analysis of 2D and 3D distance measurements to study spatial genome organization. Methods (2017).

29. V. Roukos et al., Spatial dynamics of chromosome translocations in living cells. Science 341, 660–664 (2013).

30. S. S. P. Rao, M. H. Huntley, N. C. Durand, E. K. Stamenova, A 3D Map of the Human Genome at Kilobase Resolution Reveals Principles of Chromatin Looping. Cell 159, 1665–1680 (2014).

31. P. Olivares-Chauvet et al., Capturing pairwise and multi-way chromosomal conformations using chromosomal walks. Nature 540, 296–300 (2016).

32. G. Fudenberg, M. Imakaev, C. Lu, A. Goloborodko, N. Abdennur, L. A. Mirny, Formation of Chromosomal Domains by Loop Extrusion. Cell Reports 15, 2038–2049 (2016).

33. A. L. Sanborn et al., Chromatin extrusion explains key features of loop and domain formation in wild-type and engineered genomes. Proc. Natl. Acad. Sci. U. S. A. 112 E6456–65 (2015).

34. R. A. Beagrie et al., Complex multi-enhancer contacts captured by genome architecture mapping. Nature 543, 519–524 (2017).

35. T. Sexton et al., Three-dimensional folding and functional organization principles of the Drosophila genome. Cell 148, 458–72 (2012)

36. F. Ay et al., Identifying multi-locus chromatin contacts in human cells using tethered multiple 3C. BMC Genomics 16 121 (2015)

37. M. A. Ferraiuolo et al., The three-dimensional architecture of Hox cluster silencing. Nucleic Acids Res. 38, 7472–84 (2010).

38. C. Morey, M. R. Da Silva, P. Perry, and W. A. Bickmore, Nuclear reorganisation and chromatin decondensation are conserved, but distinct, mechanisms linked to Hox gene activation. Development. 134: 909–919 (2007).

39. J. C. Fung, W. F. Marshall, A. Dernburg, D. A. Agard, J. W. Sedat, Homologous chromosome pairing in Drosophila melanogaster proceeds through multiple independent initiations. J Cell Biol 141, 5–20 (1998).

40. J. R. Chubb, S. Boyle, P. Perry, W. A. Bickmore, Chromatin motion is constrained by association with nuclear compartments in human cells. Curr. Biol. 12, 439–445 (2002).

41. W. F. Marshall et al., Interphase chromosomes undergo constrained diffusional motion in living cells. Curr. Biol. 7, 930–939 (1997).

42. R. Kalhor, H. Tjong, N. Jayathilaka, F. Alber, L. Chen, Solid-phase chromosome conformation capture for structural characterization of genome architectures. Nature Biotechnology 30, 90–98 (2012).

43. T. J. Stevens et al., 3D structures of individual mammalian genomes studied by single-cell Hi-C. Nature 544, 59–64 (2017).

44. J. A. Benanti, D. A. Galloway, Normal Human Fibroblasts Are Reisistant to RAS-Induced Senescence. Mol Cell Biol 24, 2842–52 (2004).

45. N. Naumova, et al., Organization of the mitotic chromosome. Science 342, 948–53 (2013).

46. E. P. Nora, et al., Targeted degradation of CTCF decouples local insulation of chromosome domains from genomic compartmentalization. Cell 169, 930–44 (2017).

47. M. Imakaev, et al., Iterative correction of Hi-C data reveals hallmarks of chromosome organization. Nat Methods 9, 999–1003 (2012).

48. B. R. Lajoie, J. Dekker, and N. Kaplan, The Hitchhiker’s guide to Hi-C analysis: practical guidelines. Methods 72, 65–75 (2015).

49. B. Burman, T. Misteli, G. Pegoraro, Quantitative detection of rare interphase chromosome breaks and translocations by high-throughput imaging. Genome Biology 16, 146–146 (2015).

50. S. Shachar, Ty C. Voss, G. Pegoraro, N. Sciascia, T. Misteli, Identification of Gene Positioning Factors Using High-Throughput Imaging Mapping. Cell 162, 911–923 (2015).

51. K. J. Meaburn, Fluorescence in situ hybridization on 3D cultures of tumor cells. Methods Mol Biol 659 323–36 (2010).26

52. R. C. Team, R: A Language and Environment for Statistical Computing. (2015).

53. J. French, SpatialTools: Tools for Spatial Data Analysis. R package version 1.0.2., (2015).

54. H. Wickham, The Split-Apply-Combine Strategy for Data Analysis. Journal of Statistical Software 40, 1–29 (2011).

55. H. Wickham, R. Francois, dplyr: A Grammar of Data Manipulation. R package version 0.4.3., (2015).

56. H. Wickham, ggplot2: Elegant Graphics for Data Analysis. (Springer-Verlag, New York, 2009).

57. M. Dowle et al., data.table: Extension of Data.frame. R package version 1.9.6., (2015).

58. Y. Xie, in Implementing Reprodubible Computational Research, V. Stodden, F. Leisch, R. D. Peng, Eds. (Chapman and Hall/CRC, 2014).

59. H. Wickham, stringr: Simple, Consistent Wrappers for Common String Operations. R package version 1.0.0. (2015).

60. C. A. Schneider, W. S. Rasband, K. W. Eliceiri, NIH Image to ImageJ: 25 years of image analysis. Nature Methods 9, 671–675 (2012).

